# *Bacteroides muris* sp. nov. isolated from the cecum of wild-derived house mice

**DOI:** 10.1101/2022.05.18.490454

**Authors:** Hanna Fokt, Rahul Unni, Urska Repnik, Ruth A Schmitz, Marc Bramkamp, John F. Baines, Daniel Unterweger

## Abstract

Two bacterial strains, KH365_2^T^ and KH569_7, were isolated from the cecum contents of wild-derived house mice. The strains were characterized as Gram-negative, rod-shaped, strictly anaerobic, and non-motile. Phylogenetic analysis based on 16S rRNA gene sequences revealed that both strains were most closely related to *Bacteroides uniformis* ATCC 8492^T^. Whole genome sequences of KH365_2^T^ and KH569_7 strains have a DNA G+C content of 46.02% and 46.03% mol, respectively. Most morphological and biochemical characteristics did not differ between the newly isolated strains and classified *Bacteroides* strains. However, the average nucleotide identity (ANI) and dDNA-DNA hybridization (dDDH) values clearly distinguished the two strains from described members of the genus *Bacteroides*. Here, we present the phylogeny, morphology, and physiology of a novel species of the genus *Bacteroides* and propose the name *Bacteroides muris* sp. nov., with KH365_2^T^ (DSM XXX = CCUG XXX) as type strain.

## Introduction

The mammalian intestinal tract is densely populated by microorganisms that share an ancient evolutionary history with the host (Bäckhed et al. 2005). Two dominant phyla of the intestinal microbiomes of humans and mice are the Gram-negative Bacteroidetes and Gram-positive Firmicutes (Ley et al. 2005, 2008; Reyes et al. 2010). One of the prominent genera of Bacteroidetes, *Bacteroides*, is linked to health-related traits of the host (Wang et al. 2020), provides protection against pathogens (Hecht et al. 2016; Li et al. 2017), and modulates host immunity (Round and Mazmanian 2009; Round et al. 2011). Moreover, *Bacteroides* are metabolically versatile and offer key benefits to the host, including the breakdown of dietary carbohydrates (Comstock 2009). Because of their high abundance and medical relevance, *Bacteroides* species have become important models for understanding gut microbial ecology and the role of the microbiota in health and disease (Bencivenga-Barry et al. 2020; Donaldson et al. 2020).

The first *Bacteroides* sp. was described at the end of the 19^th^ century (*Bacteroides fragilis*) (Veillon, A., Zuber 1898). Since then, 35 different *Bacteroides* species have been isolated, characterized, and validated (www.bacterio.net/bacteroides.html; García-López *et al*., 2019). A considerable number of species that were formerly classified as part of the genus *Bacteroides* were recently assigned to new genera (García-López et al. 2019). Recent microbiome studies include the genome sequence of a yet unclassified *Bacteroides* sp. (assembly accession GCA_004793475.1) that is referred to as *Bacteroides* sp. NM69_E16B (NCBI taxonomy) or *Bacteroides* sp002491635 (GTDB taxonomy) (Afrizal et al. 2022; Beresford-Jones et al. 2022; Wong et al. 2022). This genome sequence indicates the existence of an additional *Bacteroides* species that is not yet phenotypically characterized, which could be important for understanding the diversity and evolution of the *Bacteroides* genus.

Here, we describe strains KH365_2^T^ and KH569_7, which were isolated from the cecum of wild-derived house mice (*Mus musculus musculus*) at the animal facility of the Max Planck Institute for Evolutionary Biology in Plön, Germany. The two strains exhibit phenotypic characteristics of *Bacteroides* and are close relatives to the independently isolated NM69_E16B strain. All three strains are similar enough to belong to the same species and sufficiently distinct from classified *Bacteroides* species to propose *Bacteroides muris* as their species name, which was recently independently suggested on a preprint server for one of the three strains (Afrizal et al. 2022).

## Materials and methods

### Isolation and preservation

Strains KH365_2^T^ and KH569_7 were isolated from the cecal contents of two *Mus musculus musculus* house mice. The mice were wild-derived and maintained as outbred colonies in the animal facility at the Max Planck Institute for Evolutionary Biology in Plön, Germany.

Isolation and cultivation of bacteria from the mouse gut were performed in July 2018. Cecum content was collected from freshly sacrificed mice and stored at -70 °C in anaerobic Brain Heart Infusion (BHI) broth (Merck) containing 20% (v/v) glycerol. To isolate individual strains, 100 µl of the homogenized cecum content sample was serially diluted in 900 µl of anaerobic BHI medium and shortly mixed by vortexing. Next, 50 µl of the dilution was spread onto Schaedler anaerobe kanamycin vancomycin selective agar with lysed horse blood (SKV) (Thermo Fisher Scientific Inc.). After incubation for 72 h at 37 °C under anaerobic conditions, single colonies were picked and transferred onto new SKV plates. Single isolates were preserved at -70 °C in anaerobic BHI medium containing 20% glycerol. All the isolation and cultivation steps were performed under strictly anaerobic conditions in an anaerobic chamber (Coy Laboratory Products Inc.) with an atmosphere containing 5% H_2_, 5% CO_2_, and 90% N_2_. Anaerobic media were supplemented with 0.5 g/l L-cysteine hydrochloride (Sigma-Aldrich) added after autoclaving, to remove oxygen from the solution. All the materials and media were placed into the anaerobic chamber on the day prior to an experiment. The strains were submitted to the German Collection of Microorganisms and Cell Cultures (DSMZ) and to the Culture Collection University of Gothenburg (CCUG) (KH365_2^T^: DSM XXX=CCUG XXX and KH569_7: DSM XXX=CCUG XXX). The type strain *B. uniformis* ATCC 8492^T^ was obtained from the DSMZ.

### 16S rRNA gene phylogeny

Near full-length 16S rRNA gene sequences for both strains (1527 bp) were extracted from the respective whole genome sequences using the ContEst16S tool (Lee et al. 2017). The 16S rRNA gene sequences of the two novel strains were aligned against 35 *Bacteroides* type strains with validly published names (www.bacterio.net/genus/bacteroides.html) using MUSCLE (Edgar 2004). Neighbour-joining (Saitou and Nei 1987), maximum-likelihood (Felsenstein 1981) and maximum-parsimony (Kolaczkowski and Thomton 2004) trees were constructed using PAUP4 (Swofford 2002) with bootstrap analysis of 1000 random replicates and *Prevotella bryantii* as the outgroup.

### 16S rRNA gene comparison

The 16S rRNA gene sequences of strains KH365_2^T^ and KH569_7 were compared to the other closely related *Bacteroides* type strains, and the sequence similarity was determined using progressive pairwise aligner in Geneious Prime 2022.1.1 (https://www.geneious.com).

### Genome sequencing

The genomic DNA from pure cultures of strains KH365_2^T^ and KH569_7 was extracted using the DNeasy UltraClean Microbial Kit (Qiagen) following the manufacturer’s instructions. DNA samples were prepared according to the Illumina Nextera XT protocol and the final DNA library was sequenced on an Illumina NextSeq 500 system using the NextSeq 500/550 High Output Kit v2.5 with 300 cycles. Genome assemblies were generated using SPAdes 3.14 genome assembler (Bankevich et al. 2012) and annotated using Prokka (Seemann 2014).

### Genome analysis and comparison

The genomes of KH365_2^T^ and KH569_7 strains were analyzed using the Type Strain Genome Server (TYGS) (Meier-Kolthoff and Göker 2019). The pairwise digital DNA-DNA hybridization (dDDH) values between the two strains and their 19 most closely related *Bacteroides* type strains (automatically determined by TYGS) were calculated. The degree of genomic similarity between KH365_2^T^ and KH569_7 strains and closely related species was estimated using the average nucleotide identity (ANI) calculator (Yoon et al. 2017) with the species delineation threshold of < 95% for orthoANIu values. Secondary metabolite biosynthetic gene clusters were predicted using antiSMASH 6.0 (Blin et al. 2021) with the detection relaxed option.

### Microscopy

To analyze the colony morphology of the strains, they were grown on CM plates for 72 h at 37 °C, and the bright field images were captured using an Axio Zoom.V16 microscope (Zeiss) at 56× magnification. The size ranges of bacterial colonies were determined based on the measurement of five different colonies for each strain.

The cell morphology and ultrastructure were determined by conventional transmission electron microscopy (TEM). For negative staining, drops of bacterial cultures, grown for 24 h, were incubated on copper, 400 mesh, formvar- and poly-L-lysine-coated grids to allow bacteria to attach. After a short wash on several drops of dH_2_O, grids were incubated with 1% aqueous uranyl acetate <10 s and then the excess uranyl acetate was blotted and grids air-dried. For epoxy resin embedding and ultrastructural analysis on thin sections, bacterial cultures grown for 24 h were fixed by adding 2% glutaraldehyde (GA) in 200 mM HEPES, pH 7.4 at a 1:1 ratio. After 2 h of initial fixation, bacteria were pelleted, resuspended in 1% GA in 200 mM HEPES, and stored in the fixative until further processing. Cells were embedded in low-melting point agarose, post-fixed with 1% osmium tetroxide prepared in 1.5% potassium ferricyanide/dH2O for 1 h on ice, and contrasted en-bloc with 2% aqueous uranyl acetate for 2 h at room temperature. Dehydration was performed with an ethanol series 50-70-80-90-96-100-100%, each for minimum 15 min, followed by acetone, 2× 30 min. Tissue was progressively infiltrated with epoxy resin and polymerized at 70 °C for 2 days. Ultra-thin, 80-nm-thin sections were cut using a Leica UC7 ultramicrotome, deposited on copper, slot, formvar-coated grids, and contrasted with saturated aqueous uranyl acetate for 10 min, followed by 0.03% lead nitrate for 3 min. All grids were imaged in a Tecnai G2 Spirit BioTWIN transmission electron microscope (FEI, now Thermo Fisher Scientific), operated at 80 kV, and equipped with a LaB6 filament, an Eagle 4k x 4k CCD camera, and a TIA software (both FEI, now Thermo Fisher Scientific). Bacterial cell size ranges were determined based on the measurement of six different negatively stained cells of each strain, selected to represent overall size variation (from the smallest to the biggest cell).

### Physiological analyses

Gram-staining was performed using a Gram-stain kit (Carl Roth) according to manufacturer’s instructions. Cell motility was tested in semi-solid agar (0.5%) as previously described (Tittsler and Sandholzer 1936). Catalase activity was determined by adding 3% (v/v) H_2_O_2_ to fresh cells, and oxidase activity was determined using oxidase strips (Merck). Bacterial growth was evaluated on CM agar plates with different NaCl concentrations (0.5 - 5% (w/v) intervals of 0.5%), at different temperatures (18, 20, 25, 32, 37, 42, 45 and 47 °C), and in liquid CM medium at various pH (4.0 - 10, at intervals of 1.0 pH unit). The pH range of the basal medium was adjusted using the following buffers: 100 mM acetate buffer (for pH 4.0 - 5.0), 100 mM phosphate buffer (for pH 6.0 - 8.0) and 100 mM NaHCO_3_/Na_2_CO_3_ buffer (for pH 9.0 - 10.0) (Sorokin 2005). The pH values after autoclaving showed only minor changes.

### Metabolic analyses

The metabolic repertoire of KH365_2^T^, KH569_7, and *B. uniformis* ATCC 8492^T^ was studied by performing a series of assays using Biolog AN MicroPlates™ (Biolog, Hayward, CA), following the protocol provided by the manufacturer with minor modifications. The strains were grown on CM agar under anaerobic conditions at 37 °C for 5 days in order to obtain colonies of sufficient size. Individual colonies were picked and inoculated in AN Inoculating Fluid (Biolog, Hayward, CA) such that the turbidity (600 nm) of 0.05 - 0.1 was achieved. The Inoculating Fluid was then pipetted into Biolog AN MicroPlates™ (100 µl per well), and the initial turbidity (590 nm) was measured using a Spark® microplate reader (TECAN, Männedorf, Switzerland). The plates were then incubated in anaerobic jars at 37 °C for 5 days, after which the final turbidity (590 nm) was measured. The difference between the final and initial turbidity, ΔT (590 nm), was used to determine the extent to which each of the metabolites were utilized by the strains. The metabolites differentially utilized by the strains were identified by testing whether the difference in ΔT for each metabolite was significantly different between each pair of strains (KH365_2^T^ vs. *B. uniformis* ATCC 8492 ^T^, KH569_7 vs. *B. uniformis* ATCC 8492 ^T^, and KH365_2^T^ vs. KH569_7) using the Kruskal-Wallis test.

### Repositories

The GenBank/ENA/DDBJ accession numbers for the 16S rRNA gene sequences of strain KH365_2^T^ and KH569_7 are ON325392 and ON361133, respectively. The GenBank/ENA/DDBJ accessions numbers for the genome sequences of strains KH365_2^T^ and KH569_7 are XXXX and XXXX, respectively.

### Graphic illustrations

Some figures were created using BioRender.com.

## Results and discussion

### Phylogenetic analysis of 16S rRNA gene sequences

The taxonomy of the isolated strains was determined based on their 16S rRNA gene sequences after an initial taxonomic analysis of their whole genome sequences.

First, the genomes of KH365_2^T^ and KH569_7 were uploaded to the Type Strain Genome Server (TYGS) (Meier-Kolthoff and Göker 2019) to check whether the strains are classified as members of an already known bacterial species. The genome sequences of KH365_2^T^ and KH569_7 did not match any sequence in the database and were identified as members of a potentially novel species of the genus *Bacteroides*. These results were further supported by pairwise digital DNA-DNA hybridization (dDDH) values between the two strains and their 19 most closely related *Bacteroides* type strains (automatically determined by TYGS), which ranged from 21.7 to 49.2% and were all below the threshold of microbial species delineation (70%). The pairwise dDDH value between the strains KH365_2^T^ and KH569_7 was 87% (Table S1), indicating that they belonged to the same species (Meier-Kolthoff et al. 2014).

To build a phylogeny of *Bacteroides* species including the two strains KH365_2^T^ and KH569_7, 16S rRNA gene sequences were extracted and compared against those of 35 *Bacteroides* type strains with validly published names (www.bacterio.net/genus/bacteroides.html). KH365_2^T^ and KH569_7 clustered in a neighbor-joining tree within the genus *Bacteroides* (cluster represented by KH365_2^T^) (Fig. 1). Further, they formed a clade with *Bacteroides uniformis* ATCC 8492^T^ and *Bacteroides rodentium* JCM 16496^T^. These results were consistent with the topologies of maximum-likelihood and maximum-parsimony trees (Fig. S1 and S2). Pairwise sequence analysis of the full-length 16S rRNA gene sequences between the novel strains and other *Bacteroides* type strains revealed that KH365_2^T^ and KH569_7 had the highest sequence similarity to *B. uniformis* ATCC 8492^T^ (accession no. AB050110, 96.83%), followed by *B. rodentium* JCM 16496^T^ (AB547646, 96.75%) and *Bacteroides fluxus* YIT 12057^T^ (AB490802, 94.33%). These values were below the species delineation threshold of 98.7% (Goebel and Stackebrandt 1994). Moreover, 16S rRNA gene sequence similarity between strains KH365_2^T^ and KH569_7 was 99.86%, supporting the results of the whole genome analysis by TYGS that these two strains might represent the same novel *Bacteroides* taxon. A comparison between the herein described strains KH365_2^T^ and KH569_7, and the strain *Bacteroides* sp. NM69_E16B revealed a sequence identity above the species delineation threshold (99.8%), indicating that all three strains belong to the same novel taxon.

**Fig. 1.**
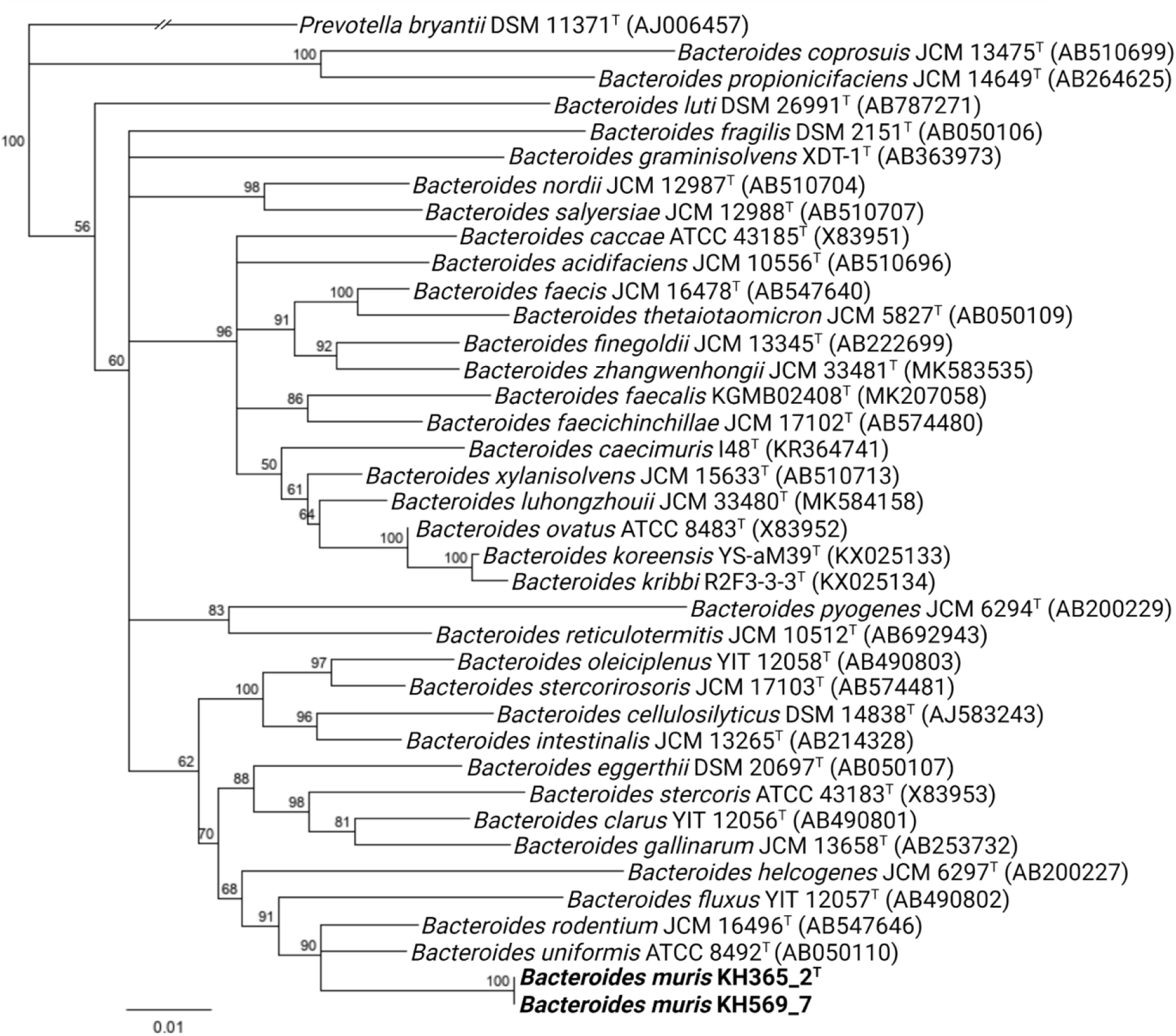
Neighbor-joining phylogenetic tree based on 16S rRNA gene sequences, showing the relatedness between *B. muris* strains KH365_2^T^ and KH569_7 (both in bold), and other members of the *Bacteroides* genus. The accession numbers of the 16S rRNA gene sequences are indicated in brackets. Numbers at nodes indicate bootstrap values (>50%) calculated from 1000 trees. *Prevoltella bryantii* DSM 11371^T^ was used as an outgroup to root the tree. The bar indicates substitutions per nucleotide position

### Genome characteristics

To study genomic features of KH365_2^T^ and KH569_7, whole genomes were sequenced and analyzed. The genomic differences between the two strains to other *Bacteroides* species, which had already been observed in 16S rRNA gene sequences (Fig. 1), were additionally determined at the level of the whole genome. The dDDH values between KH365_2^T^ and KH569_7 and their 19 most closely related *Bacteroides* type strains ranged from 21.7 to 49.2%, which is below the threshold of microbial species delineation (70%) (Meier-Kolthoff et al. 2014). The pairwise dDDH value between the two novel strains was 87% (Table S1) and thereby above the threshold. Similarly, OrthoANIu values between the novel strains and type strains of closely related species were below the species delineation thresholds, ranging from 80.07% to 80.22% similarity with *B. fluxus* YIT 12057^T^ and 92.01% to 91.85% similarity with *B. uniformis* ATCC 8492^T^ (Table S1). These results further support the assignment of KH365_2^T^ and KH569_7 to the same novel species. Moreover, the comparison of dDDH and orthoANIu confirm the classification of *Bacteroides* sp. NM69_E16B in the same species as the herein described strains (Table S2).

The draft genomes of KH365_2^T^ and KH569_7 were 4 164 660 bp and 4 177 920 bp long, with 46.02% and 46.03% mol G+C content, respectively (Table S1). Coding DNA sequences (CDSs), rRNAs, and tRNAs were annotated in the draft genomes (Table S3). A total of 3886 and 3903 CDSs were predicted for KH365_2^T^ and KH569_7, respectively. Further genome mining revealed 101 proteins involved in polysaccharide transport and degradation encoded in the genome of KH365_2^T^, while only 75 were identified in the genome of KH569_7. Moreover, several beta-lactamases were detected in the draft genomes of both strains, potentially indicating resistance to some antibiotics. The numbers of identified rRNAs and tRNAs in KH365_2^T^ were 6 and 66, respectively. Strain KH569_7 had 5 rRNAs and 48 tRNAs in its genome. These results are within the range of the number of rRNAs and tRNAs identified in closely related *Bacteroides* species (Table S3). Both strains, KH365_2^T^ and KH569_7, harbored two predicted gene clusters each for the synthesis of secondary metabolites: ribosomally synthesized and posttranslationally modified peptide classes (RiPPs) and respective recognition elements.

Taken together, whole genome sequences confirm the classification of the two strains distinct from known *Bacteroides* species, predict a function in polysaccharide metabolism among others, and reveal genomic differences between the two strains.

### Morphological characterization

To characterize the morphology of KH365_2^T^ and KH569_7, we imaged bacterial colonies and cells microscopically. The colonies of KH365_2^T^ were smooth, creamy, circular, convex, and approximately 0.8 - 1.0 mm in diameter (Fig. 2a). Their appearance was similar to colonies of *B. uniformis* ATCC 8492^T^ (Fig. 2a). The colony morphology of KH569_7 was analogous to the strain KH365_2^T^ and *B. uniformis* type strain, except for a smaller colony size (0.6 - 0.8 mm in diameter) and slightly brighter color when grown under the same conditions (Fig. 2a). The difference in colony size might be explained by differences in growth, which are also observed when growing the strains in liquid media (Fig. 2b).

**Fig. 2.**
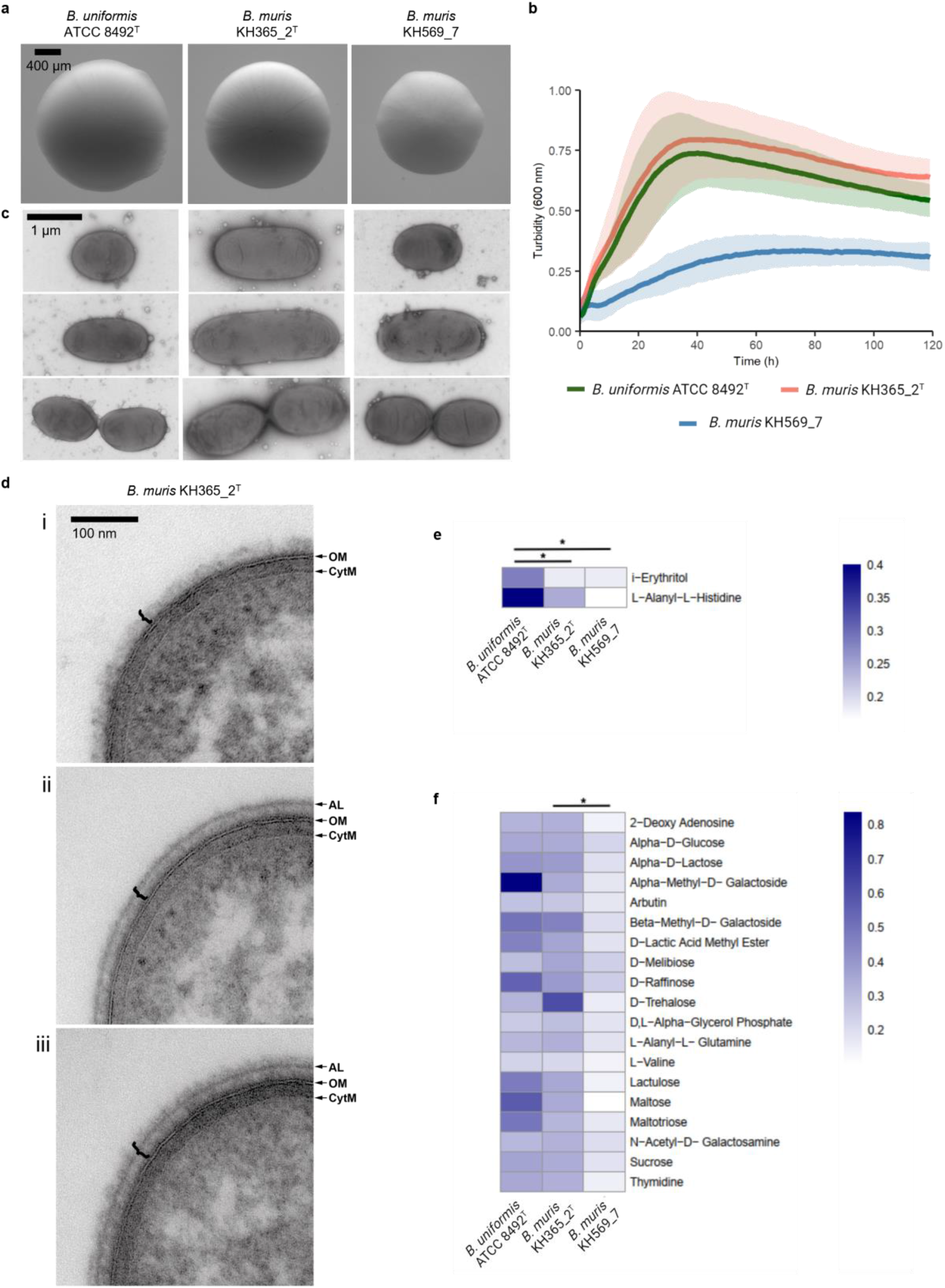
Morphology and physiology of the *B. muris* KH 365_2^T^ and KH 569_7 strains, and closely related *B. uniformis* ATCC 8492^T^. (a) Colony morphology after growth on CM modified agar for 72 hours. Magnification 56×. (b) Growth of the three strains in liquid CM modified media for 120 hours. Turbidity (600 nm) was measured every 60 min. The mean ± standard deviation of three independent experiments is shown. (c) Transmission electron microscopy images of negatively stained bacteria. (d) Ultrastructure of the cell envelope illustrating variation in the microcapsule (bracket) of *B. muris* KH365_2^T^: (i) one fringed layer, (ii) an electron dense layer with an additional, peripheral compact layer (AL), or (iii) containing three layers, including a fringed layer beyond the AL (bottom). CytM, cytoplasmic membrane; OM, outer membrane. Images were taken on transverse sections of bacteria. (e) Heatmap showing differentially metabolized compounds between *B. muris* KH 365_2^T^ and KH 569_7 strains and *B. uniformis* ATCC 8492^T^. (f) Heatmap showing differentially metabolized compounds between *B. muris* KH 365_2^T^ and *B. muris* KH 569_7 strains. The shade of color on both heatmaps represents the difference between the final and initial turbidity, ΔT (590 nm). The mean of three independent Biolog assays is shown. *: p < 0.05, Kruskal-Wallis test

Next, TEM was performed to analyze the cell morphology and ultrastructure (Fig. 2c, 2d; S3, S4). Negative staining of live bacteria that had been cultured for 24 h revealed rod-shaped cells in all three strains in the size range of 1.62 - 3.38 × 0.80 – 0.85 µm for the strain KH365_2^T^, 1.00 – 2.60 × 0.80 – 0.87 µm for the strain KH569_7, and 1.13 – 2.08 × 0.85 – 0.92 µm for *B. uniformis* ATCC 8492^T^. Notably, some *B. muris* KH365_2 cells could grow unusually long, up to 9 µm in length, which was not detected in the other two strains (Fig. S3). We observed dividing cells showing the typical constriction characteristic for the division of Gram-negative bacteria (Fig. 2c). All three strains produced outer membrane vesicles (Fig. 2c). Thin sections of resin embedded bacteria were used to characterize the ultrastructure of the cell envelope (Fig. 2d; Fig. S4). In all three strains (KH365_2^T^, KH569_7 and *B. uniformis* ATCC 8492^T^), the cell envelope consists of multiple layers: the cytoplasmic membrane, the periplasm, the outer membrane, and the microcapsule external to the outer membrane. Between independent samples of the same strain, and even within single populations, we observed bacteria with different structures of the microcapsule (Fig. 2d). This suggests that the phase variation in the capsule production observed earlier in intestinal *Bacteroides* species (Coyne and Comstock 2008) is also present in the newly described strains. The capsule could appear as i) one fringed layer (Fig. 2d i), ii) an electron dense layer with an additional, peripheral compact layer (AL, based on a description by Kornman and Holt (1981)) (Fig. 2d ii), or iii) like in (ii), but with an extra fringed layer beyond the AL (Fig. 2d iii). In KH365_2^T^ and *B. uniformis* ATCC 8492^T^, all three variations of the capsule were observed, whereas in KH569_7, only (ii) and (iii) were detected (Fig. S4).

In brief, these images show rod-shaped cells, illustrate the ultrastructure of the multi-layered cell envelope, reveal the production of outer membrane vesicles, and demonstrate variable production of the microcapsule in a single bacterial population. The two *B. muris* strains share most morphological features in common and yet differ slightly.

### Bacterial physiology

Next, we studied the physiological and biochemical characteristics of the two strains. First, their Gram-stain was assessed as a basic parameter to differentiate between bacteria. KH365_2^T^ and KH569_7 were Gram-negative (Table 1), which was to be expected based on the ultrastructure of the cell envelope (Fig. 2d) and is consistent with Gram-negative staining of *B. uniformis* ATCC 8492^T^ and other members of the genus *Bacteroides*.

**Table 1.** Characteristics of the two *B. muris* strains and the type strain of the closely related *B. uniformis* species. Strains: 1, *B. muris* KH365_2^T^; 2, *B. muris* KH569_7; 3, *B. uniformis* ATCC 8492^T^. All data from all the strains were from this study. +, positive reaction; −, negative reaction

| Characteristic | 1 | 2 | 3 |
| --- | --- | --- | --- |
| Isolation source | Murine cecum | Murine cecum | Human feces |
| pH range for growth | 5 - 10 | 5 - 10 | 5 - 10 |
| Optimum pH for growth | 7 | 6 | 7 |
| NaCl (% , w/v) range for growth | 0.5 - 5 | 0.5 - 5 | 0.5 - 5 |
| Optimum NaCl (% , w/v) for growth | 0.5 - 1 | 0.5 | 0.5 - 1 |
| Temperature range for growth (°C) | 20 - 42 | 20 - 37 | 20 - 42 |
| Optimum temperature for growth (°C) | 32 - 37 | 32 - 37 | 32 - 37 |
| Enzymatic activity: |  |  |  |
| Catalase | – | – | – |
| Oxidase | – | – | – |

Second, multiple measurements were performed to study the conditions under which the strains grow. When grown as stab cultures in aerobic conditions, KH365_2^T^, KH569_7, and *B. uniformis* ATCC 8492^T^ strains grew only at the bottom of the tube, suggesting that these bacteria are strictly anaerobic. The two strains KH365_2^T^ and KH569_7 were calatase- and oxidase-negative, as was the closely related *B. uniformis* ATCC 8492^T^ strain (Table 1). The KH365_2^T^ strain was able to grow in the presence of 0.5 - 2.0% (w/v) NaCl, similar to *B. uniformis* ATCC 8492^T^ (optimal growth at 0.5 - 1% (w/v)), whereas the KH569_7 strain could tolerate as little as 0.5% (w/v) NaCl (Table 1). Similar to *B. uniformis* ATCC 8492^T^, optimal growth of KH365_2^T^ and KH569_7 was observed at 32 - 37 °C; while KH365_2^T^ was able to grow at 20 - 42 °C and KH569_7 at 20 - 37 °C (Table 1). All three strains grew at pH 5 – 10 with slight differences in the optimum pH (Table 1).

Third, the metabolic repertoire of KH365_2^T^ and KH569_7 was systematically studied by performing a series of Biolog assays. We measured the extent to which 95 carbon sources were utilized by the novel strains and *B. uniformis* ATCC 8492^T^ (Fig. S5). Two metabolites, L-alanyl-L-histidine and i-erythritol, were differently utilized by KH365_2^T^ and KH569_7, compared to their closest relative *B. uniformis* ATCC 8492^T^, i.e. these are metabolites that are better utilized by *B. uniformis* ATCC 8492 ^T^ than by either of the novel strains (Fig. 2e). We further observed differences in the metabolism between the two strains of the novel species. Nineteen different metabolites, including lactic acid and some sugars, among others, were differentially utilized by KH365_2^T^ and KH569_7 (Fig. 2f), indicating a high degree of intra-specific diversity. Notably, all 19 of these metabolites were significantly better utilized by KH365_2^T^ than by KH569_7, and 12 of them were utilized similarly by KH365_2^T^ and *B. uniformis* ATCC 8492^T^.

Finally, we checked for motility of the strains. All were non-motile (Table 1), consistent with reports on other *Bacteroides* species (Lakhdari et al. 2011).

In summary, the physiology of the two strains is largely consistent with what might be expected from a *Bacteroides* species. Both strains lack some metabolic features of the closely related *B. uniformis* and show considerable intraspecific variation.

## Conclusion

The results of the genomic and phenotypic analysis suggest that the two herein described strains, KH365_2^T^ and KH569_7, in addition to the independently isolated strain NM69_E16B (Afrizal et al. 2022), represent a novel species of the genus *Bacteroides*, for which the name *Bacteroides muris* sp. nov. is proposed. Differences in growth, morphology, and metabolism between the strains are an indication of intraspecific diversity. This novel bacterial taxon might be present in mammals beyond mice, including humans, and holds promise to better understand the evolution of the *Bacteroides* genus as an important member of a healthy mammalian microbiota.

### Description of *Bacteroides muris* sp. nov

*Bacteroides muris* sp. nov. (mu’ris. L. gen. n. *muris* of a mouse).

Cells are Gram-negative, rod-shaped (1.62 - 3.38 × 0.80 - 0.85 µm), obligately anaerobic, non-spore-forming, and non-motile. Bacteria of this species are negative for catalase and oxidase. Colonies have a creamy, circular, and convex morphology after cultivation on CM plates at 37 °C for 72 h. Growth is observed at temperatures of 20 - 42 °C (optimum, 32 - 37 °C), at a pH of 5 - 10 (optimum, pH 7) and in presence of 0.5 - 2.0% (w/v) NaCl (optimum, 0.5%).

The type strain, KH365_2^T^, (DSM XXX = CCUG XXX) was isolated from the cecal content of a wild-derived *M. m. musculus* house mouse. This strain utilizes a wide range of carbon sources, including lactic acid. The 16S rRNA gene and genome sequences were deposited in GenBank under accession numbers ON325392 and XXX, respectively.

## Statements and Declarations

## Acknowledgements and Funding

We thank Dr. Sven Künzel for support with whole genome sequencing, Dr. Alibek Galeev for support with Gram-staining assays, Olga Vogler for help with the strain and genome submissions, the animal caretakers of the MPI for Evolutionary Biology, and Prof. Dr. Thomas Clavel for helpful comments on the manuscript. Further, we acknowledge the Central Microscopy Facility at the Department of Biology, Kiel University.

This work was funded by the Deutsche Forschungsgemeinschaft (DFG, German Research Foundation) – SFB 1182 – project-ID 261376515 to HF (Subproject B4), RAS (Subproject Z2), JFB (Subproject A2), DU (Subproject B4). RU is funded by the International Max-Planck Research School for Evolutionary Biology (IMPRS EvolBio). Work in the Unterweger Lab is supported by the German Federal Ministry for Education and Research (grant 01KI2020).

## Authors and contributors

HF, RU, and UR performed experiments, analyzed the data, and visualized the data; RAS provided resources; MB, JB and DU designed, coordinated and supervised the study. HF drafted the manuscript. All authors read, edited and approved the manuscript.

## Conflict of interest

The authors declare no conflicts of interests.

### Abbreviations

dDDH: digital DNA-DNA hybridization
ANI: average nucleotide identity
TEM: transmission electron microscopy.

