## Supplementary Information for "*Bacteroides muris* sp. nov. isolated from the cecum of wild-derived house mice"

Hanna Fokt[1], Rahul Unni[1,2], Urska Repnik[3], Ruth A Schmitz[4], Marc Bramkamp[3,4], John F. Baines[1,2], Daniel Unterweger[1,2]

[1]Max Planck Institute for Evolutionary Biology, 24306 Plön, Germany

[2]Institute for Experimental Medicine, Kiel University, 24105 Kiel, Germany

[3]Central Microscopy Facility, Kiel University, 24118 Kiel, Germany

[4]Institute for General Microbiology, Kiel University, 24118 Kiel, Germany

**Corresponding authors**

Daniel Unterweger

John F. Baines

Figure S1, S2, S3, S4, S5

Table S1, S2, S3

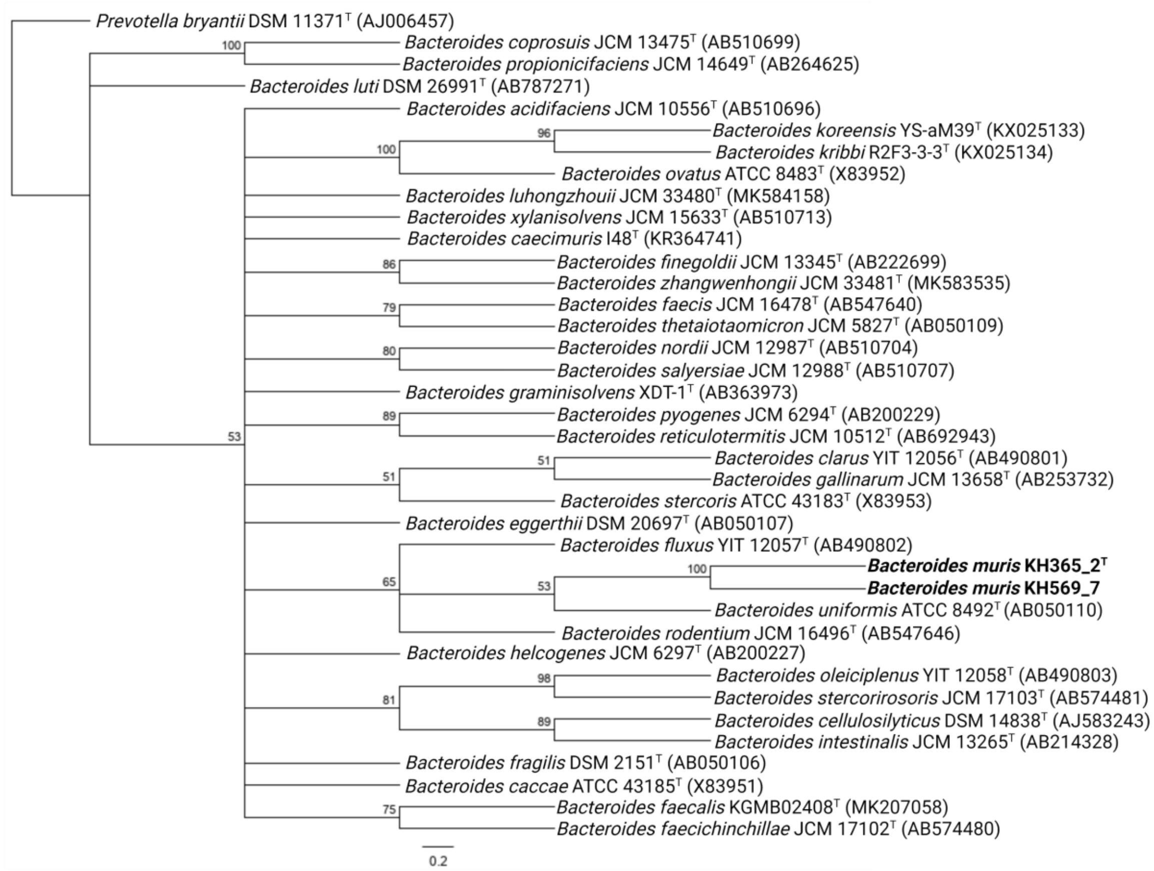


**Fig. S1** Maximum-likelihood tree based on 16S rRNA gene sequences, showing the relatedness between *B. muris* strains KH569_7 and KH365_2$^{T}$ (both in bold) and other members of the genus *Bacteroides*. The accession numbers of the 16S rRNA gene sequences are indicated in brackets. Numbers at nodes indicate bootstrap values (>50%) calculated from 1000 trees. *Prevoltella bryantii* DSM 11371$^{T}$ was used as outgroup to root the tree. The bar represents substitutions per nucleotide position.

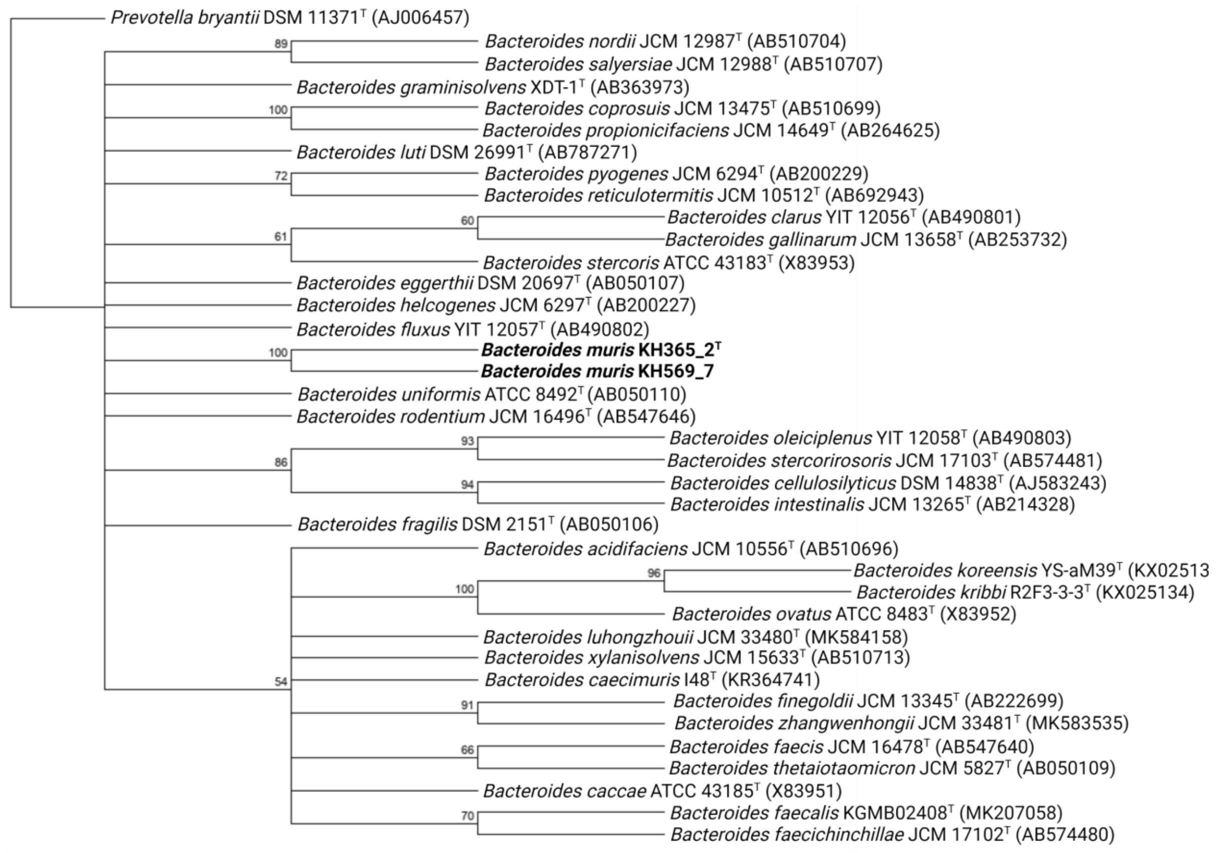


**Fig. S2** Maximum-parsimony tree based on 16S rRNA gene sequences, showing the relatedness between *B. muris* strains KH569_7 and KH365_2$^{T}$ (both in bold) and other members of the genus *Bacteroides*. The accession numbers of the 16S rRNA gene sequences are indicated in brackets. Numbers at nodes indicate bootstrap values (>50%) calculated from 1000 trees. *Prevoltella bryantii* DSM 11371$^{T}$ was used as outgroup to root the tree.

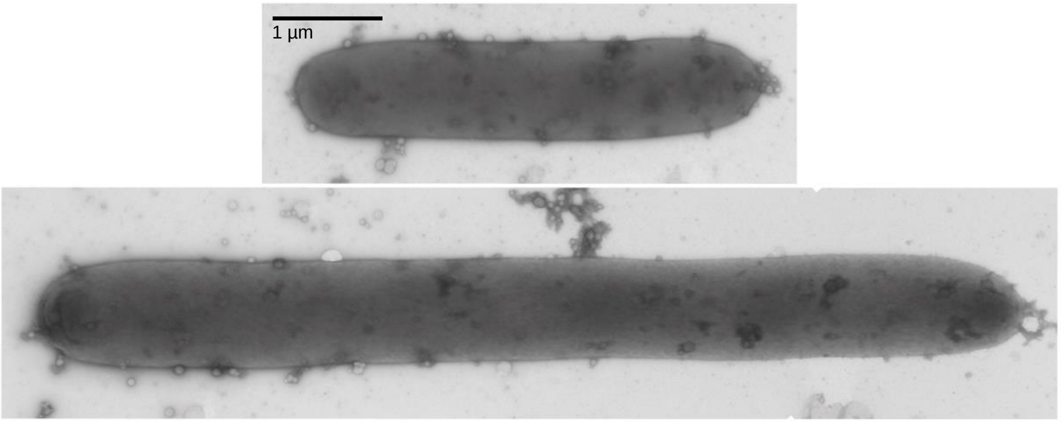


**Fig. S3** Transmission electron micrographs of negatively stained cells of *B. muris* KH 365_2$^T$. In this strain, unusually long bacteria were occasionally observed, in contrast to the other two strains.

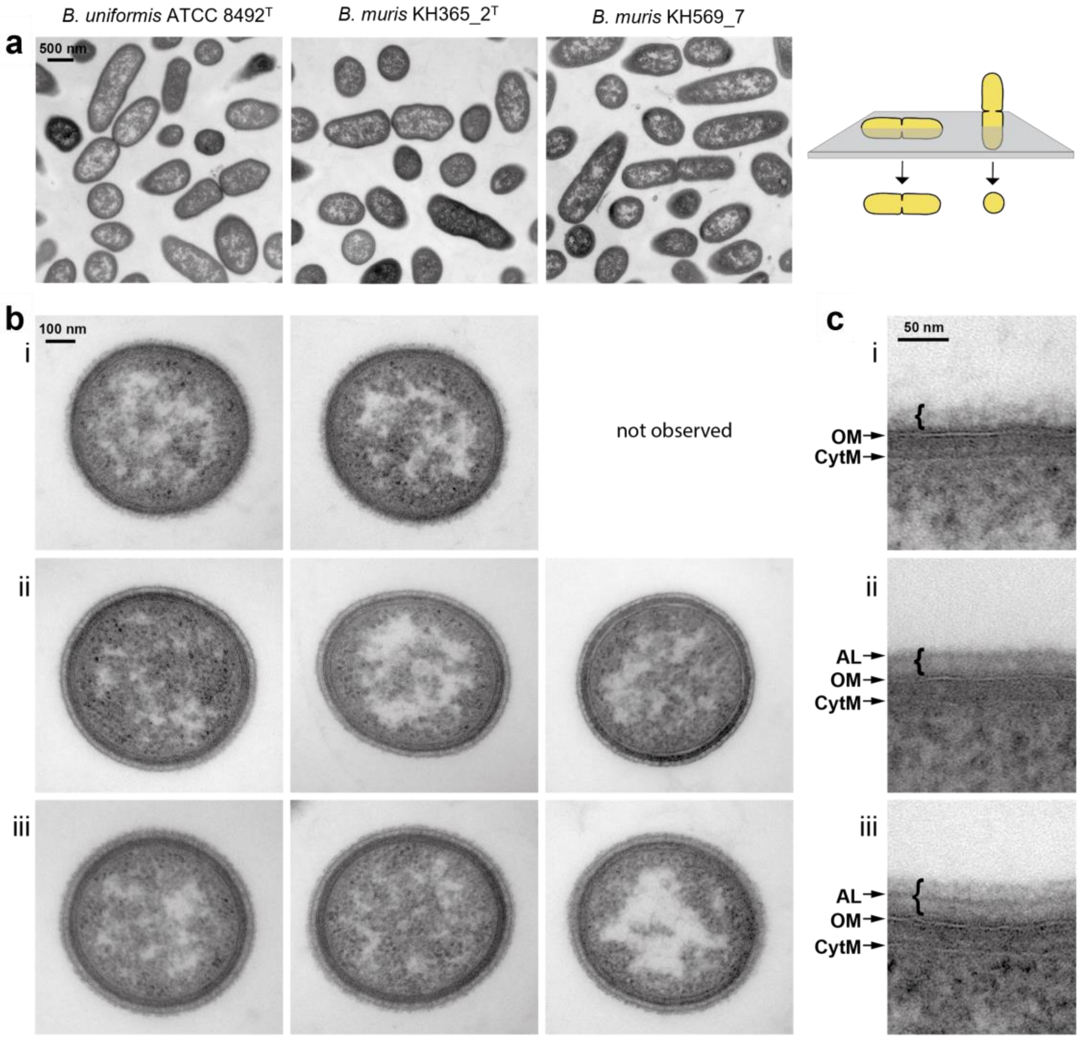


**Fig. S4** Ultrastructural analysis of *B. uniformis* ATCC 8492$^{T}$, and the newly described strains *B. muris* KH 365_2$^{T}$ and KH 569_7. Thin sections of resin embedded bacteria were imaged by transmission electron microscopy. (a) Shape of cell profiles depends on the orientation of the sectioning plane through a bacterium. Bacteria sectioned in longitudinal and in transverse orientation appear elongated and circular respectively (scheme). (b) Comparison of the structure of the cell envelope seen on transverse sections. In addition to the two bilayered membranes characteristic for Gram-negative bacteria, the capsule-like structure lies external to the outer membrane. Variation in the appearance of the capsule was observed in all three strains. (c) Detailed view of the cell envelope illustrating variation in the microcapsule (bracket): (i) one fringed layer, (ii) an electron dense layer with an additional, peripheral compact layer (AL), or (iii) containing three layers, including a fringed layer beyond the AL (bottom). CytM, cytoplasmic membrane; OM, outer membrane. All three images were taken on longitudinal sections of *B. muris* KH 365_2$^{T}$.

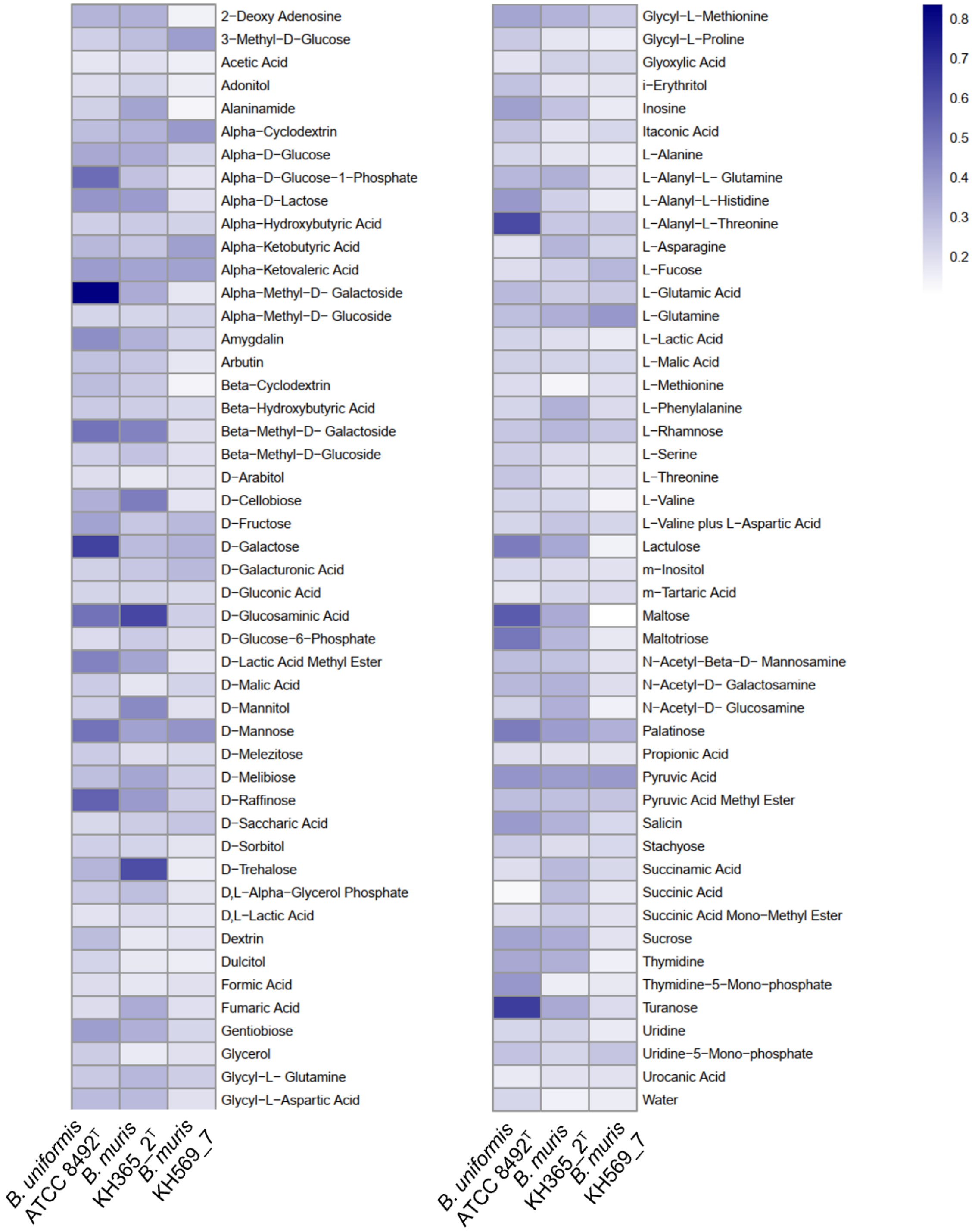


**Fig. S5** Heatmap showing the utilization patterns of different metabolites by *B. muris* KH 365_$2^{T}$, *B. muris* KH 569_7 and *B. uniformis* ATCC $8492^{T}$ strains. The shade of color on both heatmaps represents the difference between the final and initial turbidity, $\Delta T$ (590 nm) from the Biolog assay. The mean from of three independent experiments is shown.

**Table S1.** Average nucleotide identity (orthoANIu) and digital DNA-DNA hybridization (dDDH) values of *B. muris* KH365_2$^T$, *B. muris* KH569_7 and closely related *Bacteroides* strains.

| **Strains** | **Genbank accession No.** | **orthoANIu (%)** | | **dDDH (%) *** | | **G+C (% mol)** |
|---|---|---|---|---|---|---|
| | | KH365_2$^T$ | KH569_7 | KH365_2$^T$ | KH569_7 | |
| KH365_2$^T$ | TBD | – | **98.65** | – | **87.0** | 46.02 |
| KH569_7 | TBD | **98.65** | – | **87.0** | – | 46.03 |
| *B. uniformis* ATCC 8492$^T$ | GCA_000 154205 | 92.01 | 91.85 | 49.2 | 48.6 | 46.45 |
| *B. rodentium* JCM 16496$^T$ | GCA_000 614125 | 90.44 | 90.47 | 43.2 | 43.7 | 47.05 |
| *B. fluxus* YIT 12057$^T$ | GCA_000 195635 | 80.07 | 80.22 | 24.6 | 24.4 | 45.57 |

* Results are percentages based on calculations using Formula 2: The sum of all identities found in high-scoring segment pairs (HSPs) were divided by the overall HSP length. Formula 2 is independent of genome length and thus is more robust when applied to incomplete draft genomes (Meier-Kolthoff et al. 2013b).

**Table S2.** Average nucleotide identity (orthoANIu), digital DNA-DNA hybridization (dDDH) of *B. muris* KH365_2$^T$, *B. muris* KH569_7 and *Bacteroides* sp. NM69_E16B and their identity to other closely related species.

| **Strain** | **Genbank accession No.** | **orthoANIu (%)** | **dDDH (%) *** | **Related genomes (genome ID, species name, ANI [%]) **** |
|---|---|---|---|---|
| | | NM69_E16B | | |
| KH365_2$^T$ | TBD | 98.03 | 83.7 | GCF_000154205.1, *Bacteroides uniformis*, 92.44; GCF_000614125.1, *Bacteroides rodentium*, 90.86; GCF_000195635.1, *Bacteroides fluxus*, 81.67 |
| KH569_7 | TBD | 98.06 | 83.4 | GCF_000154205.1, *Bacteroides uniformis*, 92.29; GCF_000614125.1, *Bacteroides rodentium*, 90.93; GCF_000195635.1, *Bacteroides fluxus*, 81.58 |
| NM69_E16B | GCA_004793475.1 | – | – | GCF_000154205.1, *Bacteroides uniformis*, 92.15; GCF_000614125.1, *Bacteroides rodentium*, 90.91; GCF_000195635.1, *Bacteroides fluxus*, 81.55 |

* Results are percentages based on calculations using Formula 2: The sum of all identities found in high-scoring segment pairs (HSPs) were divided by the overall HSP length. Formula 2 is independent of genome length and thus is more robust when applied to incomplete draft genomes (Meier-Kolthoff et al. 2013).

** Results were obtained using the Genome Taxonomy Database Toolkit(Chaumeil et al. 2019).

**Table S3.** Comparison of the genomic features of strains KH365_2$^T$, KH569_7, and closely related species in the genus *Bacteroides*

Strains: 1, *B. muris* KH365_2$^T$; 2, *B. muris* KH569_7; 3, *B. uniformis* ATCC 8492$^T$; 4, *B. rodentium* JCM 16496$^T$; 5, *B. fluxus* YIT 12057$^T$

| | **1** | **2** | **3** | **4** | **5** |
|---|---|---|---|---|---|
| Genbank accession No. | TBD | TBD | GCA_000154205 | GCA_000614125 | GCA_000195635 |
| Number of contigs | 232 | 165 | 49 | 148 | 117 |
| Size (Mb) | 4.2 | 4.2 | 4.8 | 4.9 | 4.3 |
| CDS | 3886 | 3903 | 4663 | 4926 | 3921 |
| rRNAs | 6 | 5 | 13 | 3 | 3 |
| tRNAs | 66 | 48 | 63 | 57 | 58 |
| Other RNAs | 2 | 1 | 2 | 2 | 2 |
